# Integrating Knowledge Graphs into Machine Learning Models for Survival Prediction and Biomarker Discovery in Patients with Non–Small-Cell Lung Cancer

**DOI:** 10.1101/2024.02.29.582842

**Authors:** Chao Fang, Gustavo Alonso Arango Argoty, Ioannis Kagiampakis, Mohammad Hassan Khalid, Etai Jacob, Krishna Bulusu, Natasha Markuzon

**Affiliations:** Oncology Data Science, Oncology R&D, AstraZeneca, Waltham, MA, USA; Oncology Data Science, Oncology R&D, AstraZeneca, Gaithersburg, MD, USA; Oncology Data Science, Oncology R&D, AstraZeneca, Cambridge, UK

**Author notes:** Co-corresponding authors: Krishna Bulusu; Natasha Markuzon.

## Abstract

Survival prediction is a critical aspect of clinical study design and biomarker discovery. It is a highly complex task, given the large number of “omics” and clinical features, as well as the high degrees of freedom that drive patient survival. Prior knowledge can play a critical role in uncovering the complexity of a disease and understanding the driving factors affecting a patient’s survival. We introduce a methodology for incorporating prior knowledge into machine learning–based models for prediction of patient survival through knowledge graphs, demonstrating the advantage of such an approach for patients with non–small-cell lung cancer. Using data from patients treated with immuno-oncologic therapies in the POPLAR (NCT01903993) and OAK (NCT02008227) clinical trials, we found that the use of knowledge graphs yielded significantly improved hazard ratios, including in the POPLAR cohort, for models based on biomarker tumor mutation burden compared with those based on knowledge graphs. Use of a model-defined mutational 10-gene signature led to significant overall survival differentiation for both trials. We provide parameterized code for incorporating knowledge graphs into survival analyses for use by the wider scientific community.

## INTRODUCTION

The identification of patients who are likely to benefit from a specific therapeutic intervention is crucial to the success of clinical trials. This is particularly the case in oncology, where disease progression can be rapid and the side effects of treatment can have a significant impact on patients’ quality of life. The task of optimal patient selection is often addressed by using survival modeling analysis, in which the likelihood that a patient’s disease will respond to treatment is predicted on the basis of a combination of clinical and genomic factors. Improving patient-level survival prediction involves combining multiple modalities,^1,2^ incorporating nonlinear models,^2,3^ or performing individual biomarker hypothesis testing.^4^ Although some methods use prior knowledge, this is not done in a generalizable or replicable fashion.

Biomedical knowledge graphs (KGs) incorporate entities and their relationships in graph-based representations derived from the literature and data, including compounds, essays, genes, drug targets, protein variants, pathways, cellular components, and associated diseases. KGs have recently become prominent in biomedical and drug discovery research to extract and integrate prior knowledge.^5–7^ Their value in collecting and organizing prior biological knowledge and uncovering novel or hidden connections and the biological effects of drug responses is well documented.^6,7^

Recent studies have introduced biomedical KGs that integrate diverse public data sets to provide a single platform for answering core biomedical questions, such as disease gene identification, biomarker discovery, and drug-drug interactions.^5,7,8^ Several biologically inspired KGs are widely used, including PrimeKG,^5^ HetioNet,^7^ Orphanet,^8^ and Wikidata,^9^ among others. The Biological Insights Knowledge Graph (BIKG)^6^ is AstraZeneca’s internal KG that integrates data from more than 50 data sources, capturing more than 130 million biomedical relationships. The BIKG has been successfully used to map a prior knowledge space to focused clinical trial data to improve disease understanding and signal identification.^10^

Although various studies have explored the biomedical relationships derived from KGs,^5,10^ the application of prior knowledge from KGs in patient-level survival prediction models has not been as widely investigated. Noteworthy developments include the use of deep graph-based networks for survival prediction with breast cancer samples^11^ and a KG-based long short-term memory model to predict mortality in patients with acute kidney injury.^12^ Methods for extracting knowledge relevant to survival modeling from KGs remain an active area of research and include graph-based feature extraction,^13^ ontology-based integration,^14^ and literature-based knowledge extraction.^15^

This study aimed to incorporate prior knowledge captured in KGs to machine learning–based models for predicting patient survival and to demonstrate the improvement of these models’ predictive power, including in patients with non–small-cell lung cancer (NSCLC) treated with immuno-oncologic (IO) therapies. To this end, we developed a methodology for knowledge extraction from KGs that is suitable for patient-level survival analysis by using ML-based predictive survival models. We observed a measurable advantage of using prior knowledge in predicting the survival rate of NSCLC patients. Here we compare the performance of different KGs, including BIKG^10^ and Hetionet (heterogeneous information network),^7^ graph embedding, and modeling methods. We also provide a reusable pipeline for predicting patient survival based on combining the patient’s genomic data and the user’s choice KG. Finally, we present the biological relevance of the findings, suggesting that the proposed methodology of information extraction from KGs specific to IO treatment in patients with NSCLC can capture relevant signals out of millions of KG entities and relations.

## RESULTS

### Framework for prediction of patient survival and response and biomarker discovery

Fig. 1 depicts an overall framework incorporating KGs in the patient survival prediction model. Using the patient’s genomic signature, each gene information (e.g., mutation status) is projected to the KG, expanding knowledge to include gene-gene interactions through connected KG pathways. The identified connected graph is then projected to a lower-dimensional embedding by using the SocioDim algorithm.^16^ The gene-specific embeddings are then aggregated to a patient-level one that serves as an input for the ML patient survival prediction model. During the test phase, patient data are transformed into patient-level embedding that serves as input to a pretrained patient survival prediction model. Fig. 1b provides an enlarged view of the generation of gene-level low-dimensional embedding. For the results reported here, a 16-dimensional embedding was used.

**Fig. 1.**
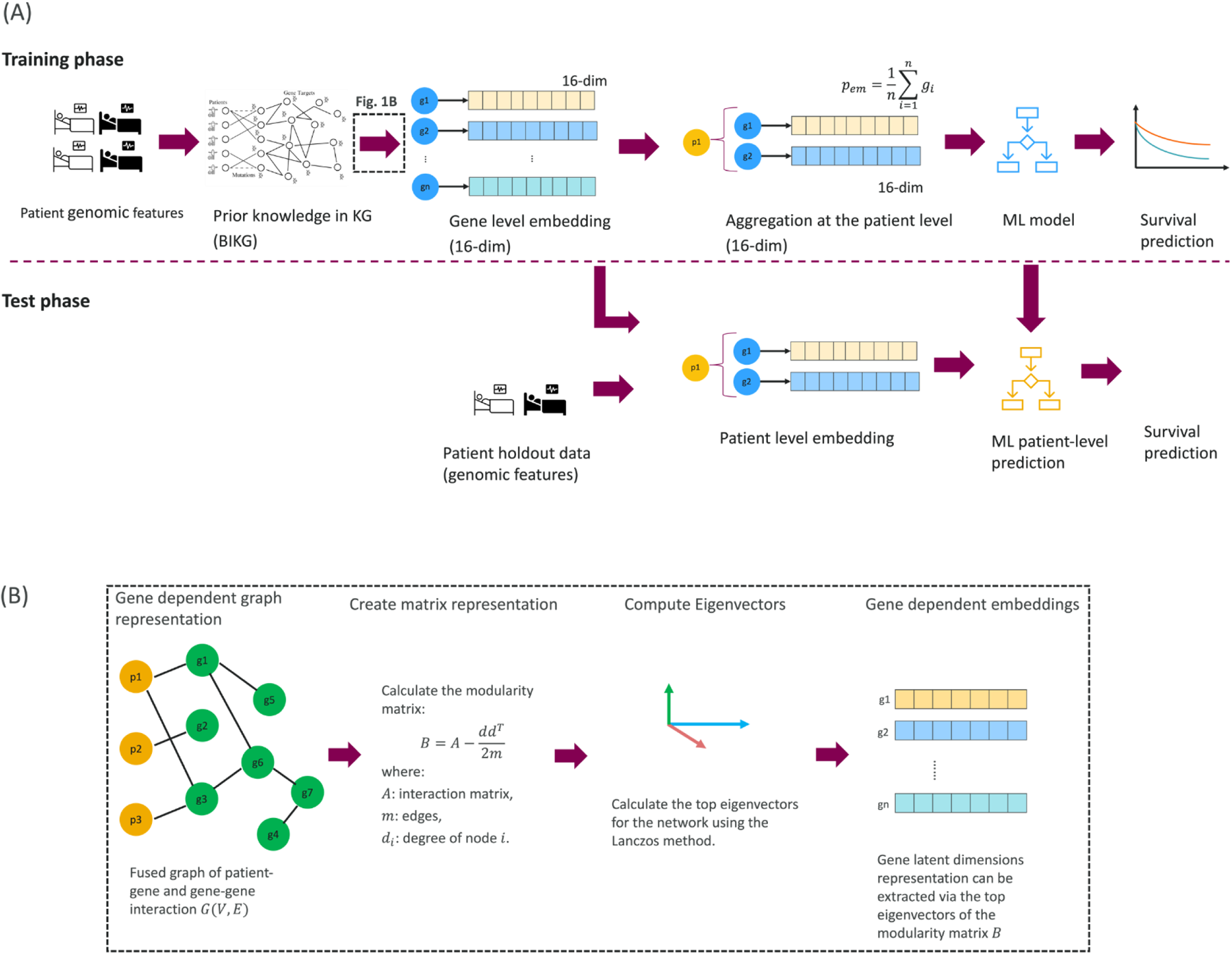
Framework incorporating KGs into ML-based models for patient survival prediction. **a**, Framework for patient survival prediction incorporating KG as input to the ML OS prediction model. **b,** Patient-level genomic data served as an input to the KG, which expands gene-gene relationships based on internal KG structure. For each gene in the panel, we identified a subgraph of interactions within the KG, which is projected to gene-specific embedding in a lower-dimensional space using the SocioDim algorithm.

We evaluated a number of other methods of generating low-dimensional embedding from KGs, including GLEE,^17^ NetMF,^18^ RandNE,^19^ NodeSketch,^20^ and BoostNE,^21^ indicating the advantage of using the SocioDim algorithm for incorporating KGs into models for prediction of OS (Extended Data Fig. 1).

We developed a stand-alone software application knowledge interface (API) that enables the incorporation of patient genomic data with relevant prior knowledge from KGs as input to the ML model to generate predictions of patient survival. Here we assess and report the results of using our internally developed BIKG^10^ as the primary KG. Subsequently, we compare its performance to that of a well-established, publicly available biomedical KG, Hetionet^7^ (Extended Data Table 1), demonstrating the adaptability of our framework to various KG platforms.

**Table 1.** Recovery of lost information from the incomplete gene panel by BIKG.

| **Input to OS prediction model** | **C-index** |  |
| --- | --- | --- |
|  | **(mean ± SD)** | ***P*** |
| Full MSK panel (481 genes) | 0.600 ± 0.025 | — |
| Full MSK panel → BIKG | 0.597 ± 0.031 | 0.815 |
| 100 random genes from MSK panel | 0.527 ± 0.016 | <0.001 |
| 100 random genes from MSK panel → BIKG | 0.598 ± 0.030 | 0.873 |

Average performance on the holdout set of OS prediction models using the C-index. Variance is estimated based on 10 independent runs. BIKG is successful in recovering the loss of information from an incomplete gene panel. The input to the OS prediction model using BIKG based on an incomplete panel is as successful as input based on a full panel.

In addition, we evaluated multiple ML-based predictive algorithms for patient survival. Using random survival forest proved to be the fastest, most stable, and most generalizable method across the suite of standard methods that can use censored data, though the advantage was insignificant.

Finally, we applied the proposed framework to analyze improvements in the use of KGs to predict survival in NSCLC patients from different studies and clinical trials.

### Use of prior knowledge from KGs in OS prediction: NSCLC patient cohort

Genomic panels play a significant role in oncology to assess treatment response in patients. These panels comprise genes carefully selected through comprehensive studies and expert curation.^22^

In this first study, we assessed the predictive power for OS outcome in NSCLC patients, using a gene panel alone and in combination with prior knowledge derived from the BIKG with the MSK (Memorial Sloan Kettering)–MET (Metastatic Events and Tropisms) 2021 NSCLC primary tissue sample data set (see Methods). Although adding the BIKG to a full gene panel did not improve OS predictions, adding it to an incomplete panel (with a significant drop in predictive performance) demonstrated an ability to recover missing information and bring OS predictions to the level achieved using the entire panel. In particular, we evaluated four inputs for the OS prediction model: (a) a complete MSK gene panel (481 genes), (b) a complete MSK gene panel combined with the BIKG, (c) a random subset of an MSK gene panel (100 genes), and (d) a random subset of an MSK gene panel combined with the BIKG.

Table 1 shows a comparison of the OS predictive performance of the four above-mentioned models, using concordance index (C-index) and variance derived from 10 separate runs. The results are reported on the independent test sets. There was no statistically significant improvement in OS prediction between scenarios (a) and (b) across 10 independent runs (Table 1). When the predictive performance of the model based on a random subset of genes was compared with that based on a full panel (scenarios a and c), there was a statistically significant decrease in predictive performance (Table 1). Adding the BIKG to a reduced gene subset (scenario d) brought the performance back to that of the full panel (Table 1).

Fig. 2 shows a comparison of the models’ performance using Kaplan–Meier (KM) survival analysis.^23^ The OS prediction was used to stratify patients into high- and low-risk subgroups with a 75th percentile cutoff. The model did not effectively stratify patients when a complete gene panel (Fig. 2a) was used. In contrast, using the full gene panel combined with the BIKG achieved statistically significant patient stratification (Fig. 2b). Similarly, use of an incomplete gene panel did not result in successful stratification (Fig. 2c), whereas adding the BIKG to the input yielded statistically significant stratification (Fig. 2d). No significant difference in OS prediction was seen when primary and metastatic tissue site data were compared (Extended Data Fig. 2).

**Fig. 2.**
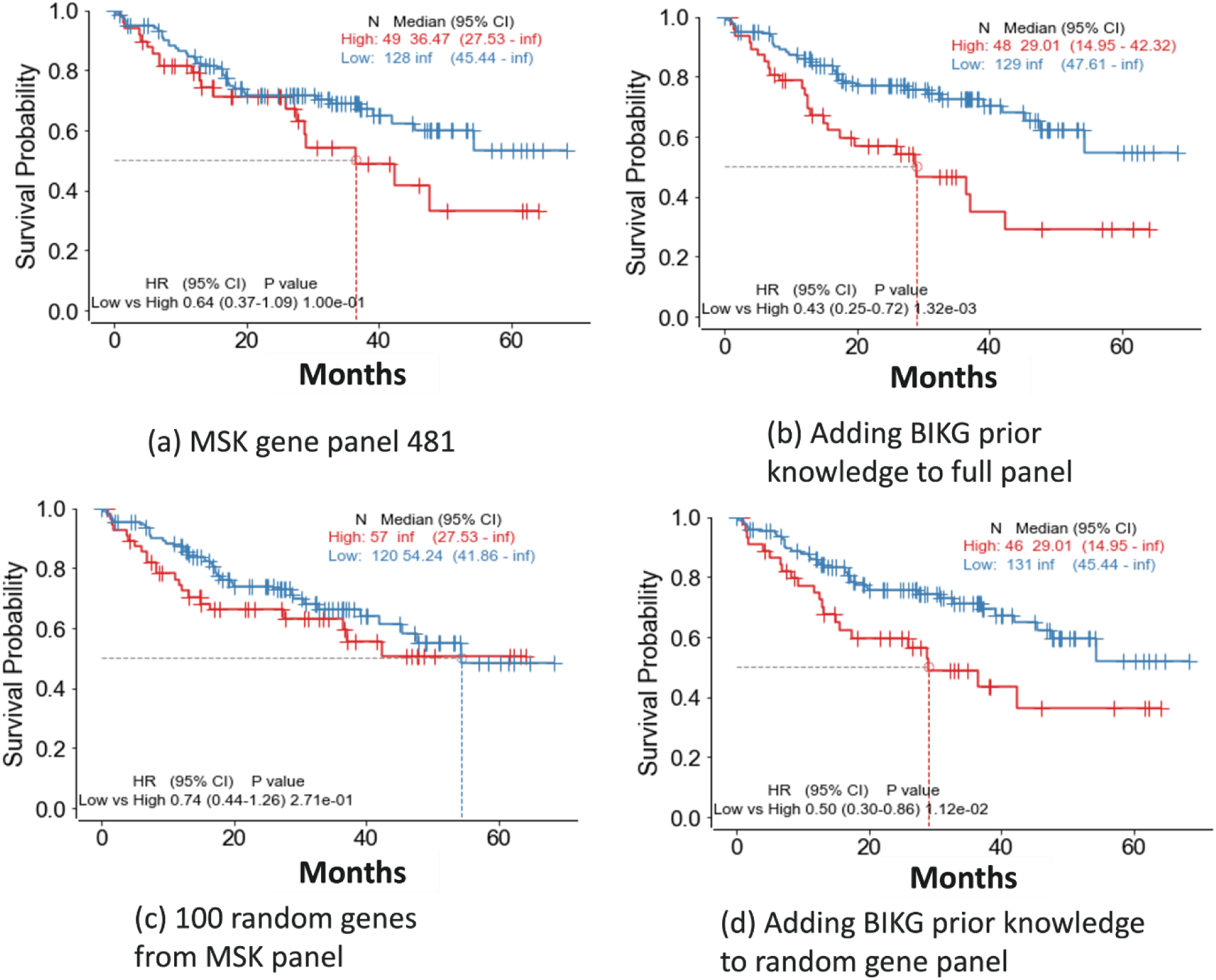
Patient OS stratification using different inputs to OS prediction models. **a**, Complete MSK gene panel; **b**, BIKG using full MSK panel; **c**, random 100 genes from the MSK panel; **d**, BIKG using a random gene panel. KM plots show statistically significant survival stratification for **b** and **d**, indicating that BIKG improved patient stratification when based on incomplete panel data, thereby complementing the incomplete MSK gene panel.

### Addition of BIKG for OS prediction in NSCLC IO-treated patients in clinical trials: POPLAR and OAK trials

The accuracy of OS prediction was improved through the incorporation of prior knowledge from the BIKG, as opposed to exclusively relying on gene panel data when analyzing two distinct cohorts of patients with late-stage NSCLC treated with IO from two separate clinical trials, OAK and POPLAR.

#### BIKG-based prediction of OS in OAK clinical trial data

We used the OAK NSCLC phase 4 gene panel data independently and in conjunction with the BIKG as input features for ML models aimed at predicting OS. Fig. 3a–c and Table 2 show independent test set results of patient stratification based on the models’ predictions using a 75% cutoff. The model trained with the BIKG in combination with the OAK gene panel data performed statistically better than the model based on the OAK gene panel only and better than that using known blood-based tumor mutation burden (bTMB) with a cutoff of 16 (Table 2).^24^

**Fig. 3.**
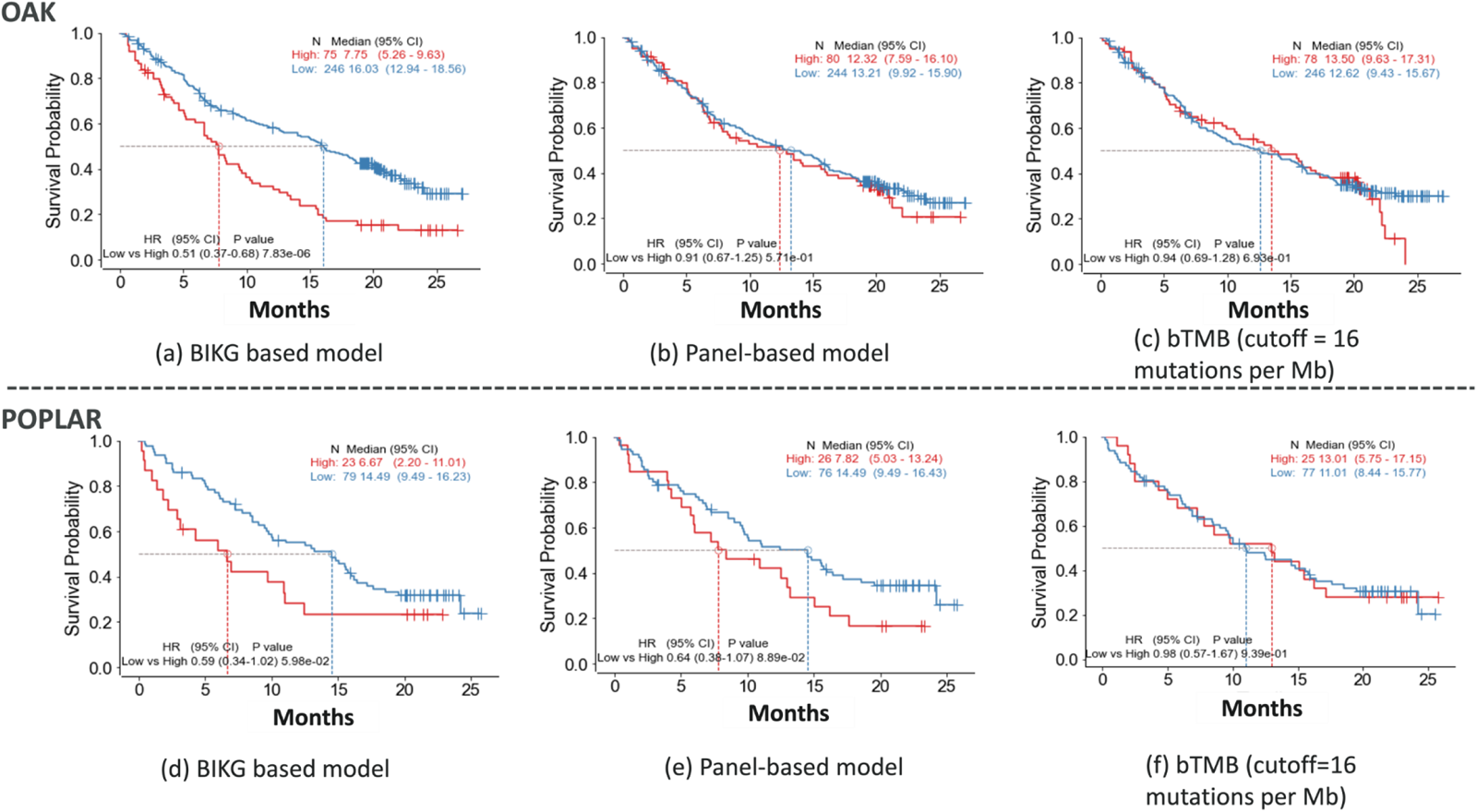
BIKG-based OS predictive performance for NSCLC IO OAK and POPLAR clinical trials. Trained using OAK, evaluation results are reported on an independent test set from the OAK clinical trial and all POPLAR clinical trial data. **a**–**c**, Evaluation of model performance using KM stratification with fivefold cross-validation on OAK data set. **a**, Model trained with BIKG using OAK gene panel data; **b**, model trained with OAK gene panel; **c**, model trained using bTMB with a cutoff of 16. The model trained with BIKG (**a**) had a statistically significant *P* value, whereas models trained with the OAK gene panel only (**b**) or bTMB only (**c**) did not. **d**–**f**, Performance on an independent POPLAR data set by models trained with the OAK data set (same models as in **a**–**c**). **d**, Model trained with BIKG in combination with the OAK gene panel; **e**, model trained with the OAK gene panel; **f**, model using bTMB with a cutoff of 16. The model using BIKG (**d**) had a statistically significant *P* value, whereas models trained with the OAK gene panel only (**e**) or bTMB only (**f**) did not show statistical significance.

**Table 2.**
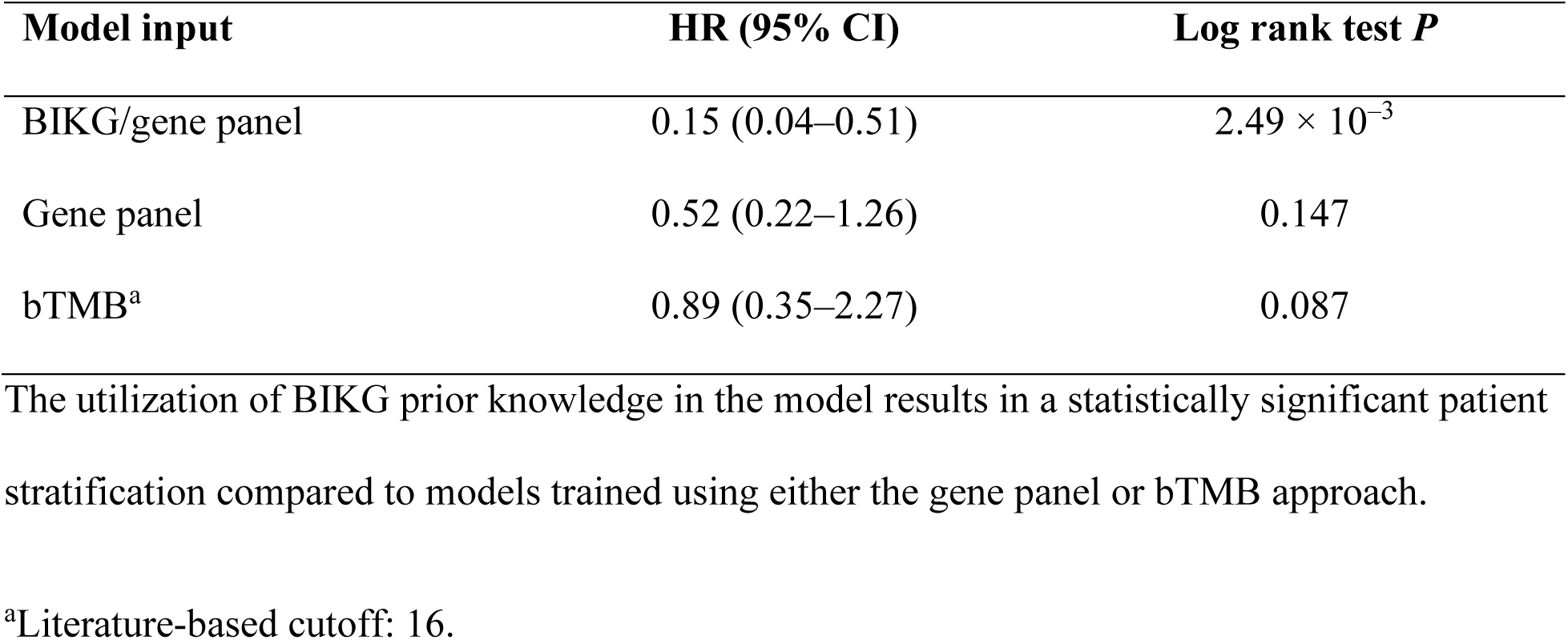
Patient OS stratification using different model inputs: OAK independent test set.

| Model input | HR (95% CI) | Log rank test *P* |
| --- | --- | --- |
| BIKG/gene panel | 0.15 (0.04–0.51) | $2.49 \times 10^{-3}$ |
| Gene panel | 0.52 (0.22–1.26) | 0.147 |
| bTMB<sup>a</sup> | 0.89 (0.35–2.27) | 0.087 |

The utilization of BIKG prior knowledge in the model results in a statistically significant patient stratification compared to models trained using either the gene panel or bTMB approach.
<sup>a</sup>Literature-based cutoff: 16.

#### BIKG-based prediction of OS in POPLAR clinical trial data

The developed OAK-based models were further evaluated by using data from an independent phase 2 clinical trial, POPLAR^25^ (Fig. 3d–f, Table 3). Similar to the evaluation using an independent test set from the OAK trial, a model trained with the BIKG in combination with the OAK gene panel performed statistically better than the model based on the OAK gene panel only. Tumor mutation burden (TMB)–based stratification with a cutoff of 16 was not as effective (Table 3).

**Table 3.** Patient OS stratification using different model inputs: POPLAR data set.

| **Model input** | **HR (95% CI)** | **Log rank test *P*** |
| --- | --- | --- |
| BIKG/gene panel | 0.59 (0.34–1.02) | $5.98 \times 10^{-2}$ |
| Gene panel | 0.64 (0.38–1.07) | 0.09 |
| bTMB<sup>a</sup> | 0.98 (0.57–1.67) | 0.939 |

The utilization of BIKG prior knowledge in the model results in a statistically significant patient stratification compared to models trained using either the gene panel or bTMB approach.
<sup>a</sup>Literature-based cutoff: 16.

#### Stability and performance gain of BIKG-based OS prediction models

Fig. 4 illustrates the performance of the BIKG-based model in predicting OS with the MSK and POPLAR data while evaluating models’ stability and benefit of adding BIKG for high risk versus low risk groups. The average performance indicated a consistent advantage of the BIKG-based models in predicting OS in a group of high risk patients.

**Fig. 4.**
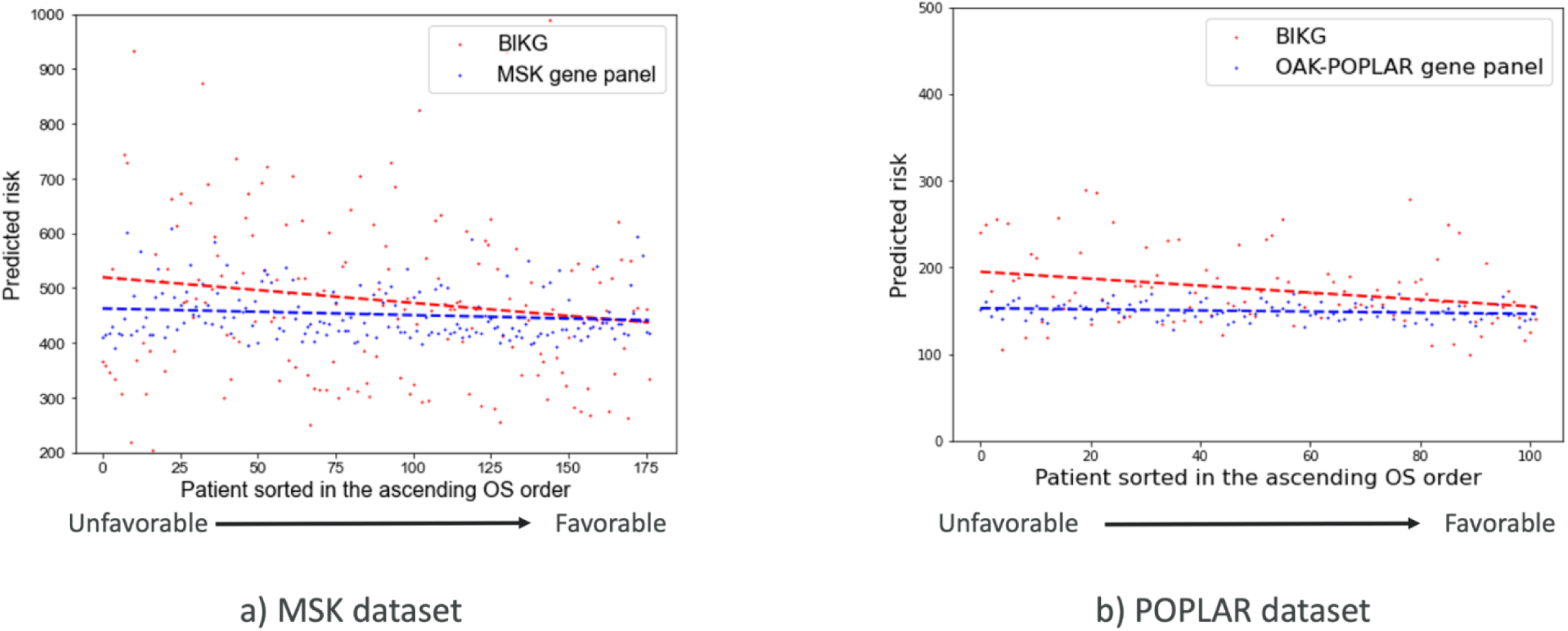
OS prediction in models using BIKG versus gene panel only. Patient survival prediction using **a**, the MSK independent test set and **b**, the POPLAR data set. Patients are sorted in ascending order by their OS. The BIKG-based model (red dashed line) indicated a much more elevated predicted risk for patients with low OS than the model based on the gene panel only (blue dashed line). The slope difference between the red and blue lines is (–0.469) – (–0.112) = (–0.347) in **a** and (–0.400) – (–0.065) = (–0.335) in **b**.

The difference in OS prediction performance between the models was much greater for patients with high predicted risk than for those with low predicted risk. Specifically, when focusing on the top 20% of patients with an unfavorable risk profile, the BIKG-based model exhibited an average predicted risk that was 15.6% higher than that of the panel-based model using the MSK data set, and 28.1% greater than that of the panel-based model using the POPLAR data set. When focusing on the top 20% of patients with a favorable predicted outcome, we noted that the BIKG-based model exhibits an average predicted risk of the BIKG-based model was only 0.5% higher than that of the model using the MSK data set, and 6% higher than the panel-based model using the POPLAR data set.

### Biomarker-based signature for OS differentiation

To identify genomic signatures associated with high and low OS, we assessed the importance of individual genes identified by the predictive model, followed by an analysis of the mutation signature of a combination of genes.

#### Importance of individual genes

Table 4 and Table 5 show the top most predictive genes identified by the developed OS prediction models across 10 independent runs.

**Table 4.** Top 10 genes identified as important in predicting OS in the OAK and POPLAR studies with BIKG-based models.

| Model-defined top<br>10 genes (no. of<br>times out of 10<br>runs)<sup>a</sup> | OAK |  | POPLAR |  |
| --- | --- | --- | --- | --- |
|  | Mutation frequency (%) | HR (95% CI) | Mutation frequency (%) | HR (95% CI) |
|  |  | using a single gene |  | using a single gene |
| *TP53* (10) | 47.83 | 1.51 (1.15–1.98) | 53.92 | 1.36 (0.84–2.20) |
| *EGFR* (9) | 9.25 | 1.29 (0.82–2.02) | 10.78 | 1.00 (0.48–2.09) |
| *ATM* (9) | 8.64 | 0.70 (0.43–1.16) | 14.71 | 0.54 (0.26–1.13) |
| *KRAS* (7) | 9.25 | 1.25 (0.80–1.96) | 7.84 | 1.00 (0.43–2.32) |
| *EPHA7* (7) | 4.32 | 1.57 (0.88–2.82) | 5.88 | 1.23 (0.49–3.06) |
| *CTNNB1* (6) | 2.16 | 1.03 (0.38–2.78) | N/A | N/A |
| *PRKDC* (6) | 8.02 | 1.47 (0.92–2.32) | 5.88 | 0.80 (0.29–2.19) |
| *NFE2L2* (6) | 5.56 | 1.16 (0.66–2.03) | 10.78 | 1.98 (0.98–4.01) |
| *SOX9* (6) | 7.09 | 1.05 (0.63–1.76) | N/A | N/A |

N/A, not applicable.

**Table 5.** Top 10 genes identified as important in predicting OS in the MSK-MET 2021 study.

| **Model-defined top 10 genes**<br>**(no. of # times out of 10 runs)<sup>a</sup>** | **Mutation frequency (%)** |  |
| --- | --- | --- |
|  | **Training data** | **Independent test data** |
|  | **(*n* = 1,602)** | **(*n* = 177)** |
| *STK11* (10) | 14.23 | 14.68 |
| *KRAS* (10) | 36.45 | 34.46 |
| *KEAP1* (10) | 13.48 | 16.94 |
| *TP53* (10) | 43.19 | 42.93 |
| *SMARCA4* (10) | 7.62 | 6.21 |
| *ATM* (10) | 7.30 | 4.51 |
| *PTPRD* (10) | 8.98 | 7.34 |
| *EPHA3* (10) | 5.49 | 7.91 |
| *RBM10* (10) | 9.80 | 10.16 |
| *FAT1* (6) | 6.86 | 6.21 |

Recent studies have highlighted the importance of *TP53*-*ATM* co-mutations as a biomarker for immune checkpoint inhibitor (ICI) response in NSCLC patients.^26,27^ Chen et al. showed that PD-L1 expression was not only high in a co-mutated cohort but was also associated with better OS than both single-gene mutations and wild-type subpopulations.^26^ Zhang et al. reported the remarkable concurrency of *TP53*, *STK11*, *PTPRD*, *RBM10*, and *ATM* mutations in patients with *KRAS*-mutant NSCLC and highlighted the significance of this relationship in assessing ICI efficacy.^28^ Bai et al. went one step further and recommended an eight-gene mutation signature to predict the response of ICI in patients with non-squamous NSCLC.^29^

#### Gene mutation signature as an OS differentiator

Individual genes identified as important by the models did not show stratification power. However, a combination of key genes identified by the BIKG-based model (*EGFR*, *TP53*, *ATM*, *PRKDC*, *STAT3*, *CTNNB1*, *KRAS*, *NFE2L2*, *EPHA7*, and *SOX9*) was used to differentiate OS in OAK and POPLAR patient cohorts (Fig. 5a, b). We considered the biomarker signature to be positive (defined in Fig. 5 as “Mut”) if there were at least two mutations among the selected genes and negative (defined in Fig. 5 as “Wt”, wild type) otherwise. Using key genes identified by the BIKG-based model, we observed a statistically significant difference in OS for the OAK data set (Fig. 5a). A similar, but not statistically significant, pattern was observed for POPLAR (potentially due to a too-small patient cohort; *n* = 57) (Fig. 5b).

**Fig. 5.**
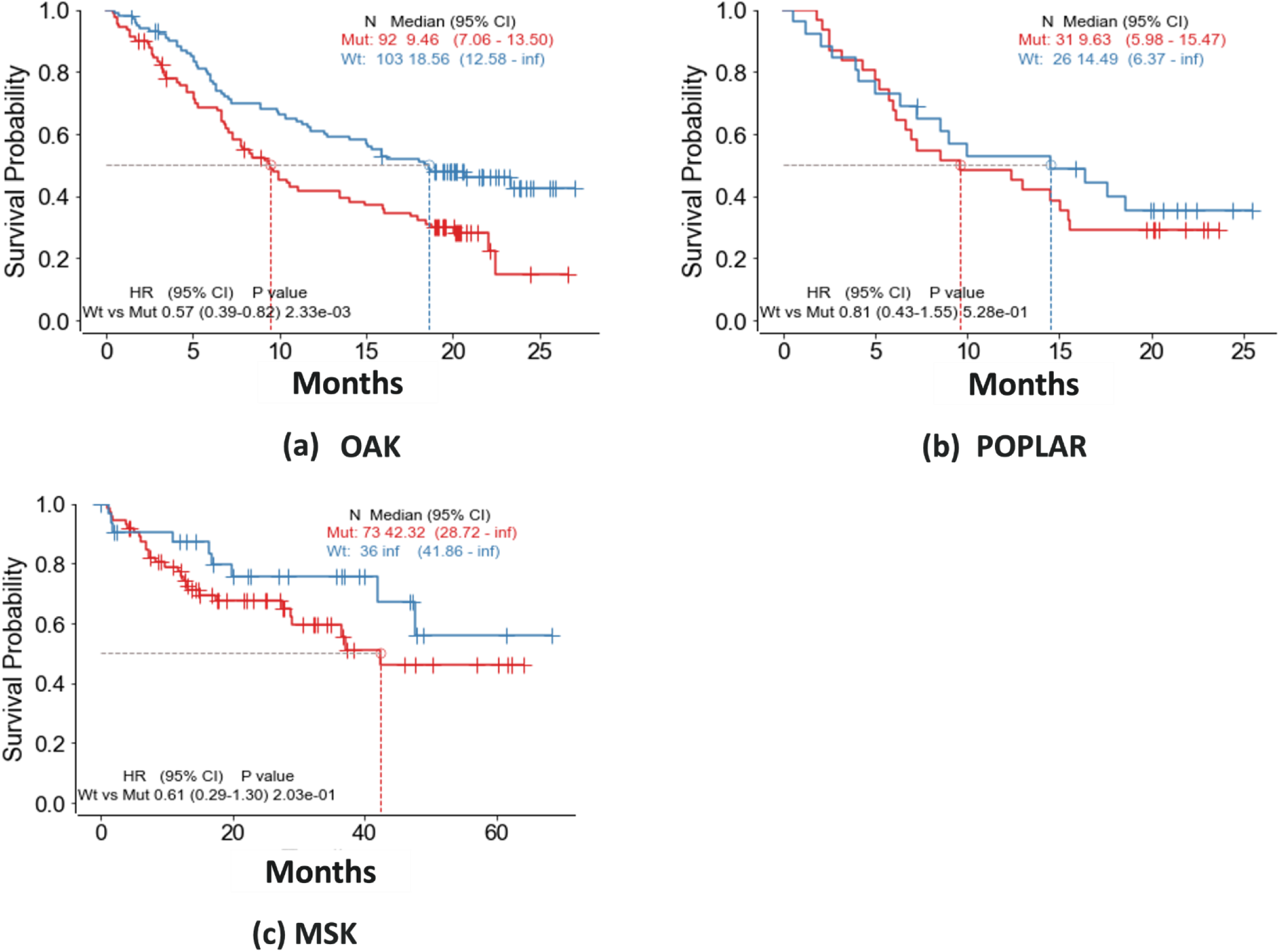
Differentiation by OS on OAK, POPLAR, and MSK data sets using mutation signature. Patient survival prediction using a, the OAK independent test set, b, the POPLAR data set, and c, the MSK independent test set. Mut, mutation biomarker; Wt, wild type.

Fig. 5c shows the results of using the gene signature of key genes (*STK11*, *KRAS*, *KEAP1*, *TP53*, *SMARCA4*, *ATM*, *EPHA3*, *PTPRD*, *RBM10*, and *FAT1*) to differentiate OS in the MSK cohort.

#### Incidence of mutations in predicted high- and low-risk groups

We compared the difference in mutation prevalence between predicted high- and low-risk groups when using the gene panel only versus the BIKG as input to the predictive survival model for the OAK and POPLAR data sets (Extended Data Table 4). Models using the BIKG identified more patients with high risk associated with *TP53*, *EGFR*, *EPHA7*, *PRKDC*, and *NFE2L2* gene mutations than low-risk patients. The high-risk patients had lower TMB on average. Similarly, we compared the high- and low-risk groups using the MSK data set (Extended Data Table 5). Models using the BIKG identified more patients with high risk associated with *STK11*, *KRAS*, *KEAP1*, and *TP53* gene mutation than low-risk patients.

## DISCUSSION

The results of this study can provide a blueprint for how integrative analytics using prior knowledge feature spaces can add value to critical clinical decisions, helping to identify the right drug for the right patient. The proposed methodology of incorporating prior knowledge into patient-level survival analysis demonstrates the advantages of automated information extraction from KGs, allowing it to speed up and boost the performance of predictive models.

### Incorporation of KG information into patient-level predictive survival models

Our methodology aims to automatically incorporate prior knowledge extracted from KGs into patient-level predictive models, specifically, ML-based patient survival prediction models. Earlier studies on core biomedical questions have not addressed the extraction of information from KGs that describe patients, concentrating more on biomedical interactions and the effects of individual genes or pathways. This study demonstrated the value of using KGs in addressing patient-level problems.

The patient survival prediction pipeline, using patient genomic features as input and prediction of patient survival as output, is a stand-alone API bringing KGs as input to patient survival prediction models. The API can incorporate different KGs and modeling methods in addition to those introduced in this study. It provides training, testing, and validation schema with added bootstrapping capability to ensure the generalization of results to previously unseen data.

Several methods of translating graph representation into low-dimensional embedding suitable for ML models have been evaluated in this study. Of those, SocioDim has demonstrated a stable advantage in improving the predictive power of patient survival models.

Comparing the BIKG and Hetionet as examples of KGs in application to NSCLC, we observed similar predictive performance of the models, with a slight advantage of BIKG. The use of disease- and treatment-specific subgraphs rather than a full graph (with BIKG) improved survival prediction, suggesting that targeting more relevant information in KGs helps to reduce the noise and uncertainty of information extracted from KGs.

Subgraphs from BIKG capture the complex biomedical relationships within a network architecture and may have the potential to provide explainable insights into disease biology and drug response. Although the performance of predictive models based on full BIKG versus selected subgraphs was not notably different, using subgraphs may allow better identification of potential biomarkers.

Of the predictive algorithms, random survival forest proved to be the most stable and generalizable across the suite of standard methods that use censored data, though the advantage was insignificant.

### Application of KG-based models for OS prediction to NSCLC patients

We observed a consistent advantage of using prior knowledge extracted from KGs for OS prediction with MSK-MET 2021, POPLAR, and OAK data in NSCLC patients. For MSK-MET 2021 data, adding BIKG to the full panel as input did not confer a significant advantage. When BIKG was added to the incomplete panel, however, we observed a remarkably automated retrieval of the missing information, allowing the predictive performance to return to the levels obtained with the complete panel. This finding suggests that BIKG can restore incomplete, missing data in a manner unbiased by human error in a fast and automated way. Identifying important genes constituting a comprehensive gene panel is a time-consuming manual process compared with the quick, automatic extraction of information from the BIKG. The low variance in performance between different bootstraps supports the superiority of the BIKG’s performance despite the graph’s high dimensionality and noisiness.

For POPLAR and OAK clinical trial data from IO-treated NSCLC patients, the performance of OS prediction was consistently improved when the BIKG was used as input than when models based on the gene panel alone or bTMB were used. The results were generalizable to the unseen data; the BIKG-based model developed with OAK clinical trial data was evaluated using POPLAR data and demonstrated consistently better performance as compared to gene panel or bTMB. As with the MSK-MET 2021 data, we observed low variance in performance between models using different bootstraps, supporting the BIKG-based model improved robustness claim.

KG-based models showed a higher gain in performance improvement for patients in the high-risk group, observed for both MSK, and POPLAR/OAK datasets, suggesting that the additional knowledge extracted from KGs contributes to the stabilization and improvement of high-risk predictions while the smaller incidence of important for the prediction model mutations in the low-risk group indeed makes KG contribution to the model less significant.

The genes that contributed most to improving OS prediction showed consistency across the models (Tables 4, 5). The co-occurrence of mutations confirmed previously observed associations (Extended Data Fig. 3). The genes identified as important by the models have strong clinical relevance from previously published studies. A complete OncoKB^30^ assessment of these genes is provided in Extended Data Tables 2 and 3. For both MSK and POPLAR/OAK datasets, BIKG-based models were able to improve the differentiation in key mutation incidence between high- and low-risk patient groups.

The identified mutational signature built from a combination of selected genes identified by predictive models proved to be an OS differentiator. For the POPLAR/OAK IO-treated population, the mutational signature showed a statistically significant difference in survival between signature-positive and signature-negative patients in the OAK cohort, with similar, but not statistically significant, results in the POPLAR cohort (Fig. 5a-b). A similar statistically significant differentiation was observed for MSK (Fig. 5c). The finding supports the importance of combining genes as a potential biomarker to amplify a patient’s response to treatment.

### Limitations and next steps

The analysis of NSCLC data was performed by using gene panel data only, limiting our ability to identify novel biomarkers. Clinical gene panels capture only a tiny fraction of relevant disease biology, presenting a significant hindrance to clinical translation of any predictive analysis. Next steps should include identifying suitable complete genome data and applying the proposed methodology to the identification of new biomarkers.

Although this study highlights the value of biomedical KGs in deciphering signals of clinical response, the biomedical and data science communities need to address critical gaps and learnings to deliver stable, reliable and scalable value. Patient survival and clinical response have multiple independent confounding factors (e.g., clinical and socioeconomic), not all of which are currently captured in a comprehensive fashion. Clinical trials capture an inherent bias in patient selection that is not reflected in existing predictive models. KGs should evolve to include nonscientific metadata to address this issue. KGs are also inherently noisy, and careful consideration is required to define context-specific graphs to address clinical questions. From a technical perspective, ML models are “data hungry,” and better models that can handle small patient cohorts need to be developed, especially given that the design of clinical trial arms is becoming increasingly specific in the era of precision medicine.

## METHODS

### Data

#### MSKCC metastatic events and tropisms

The MSK Cancer Center (MSKCC) cohort is a pan-cancer cohort of 25,775 patients with metastatic diseases across 50 tumor types.^31^ All samples were profiled by using the MSK-IMPACT (Integrated Mutation Profiling of Actionable Cancer Targets) targeted panel, which can detect mutations and other critical changes in the genes of both rare and common cancers. The data set comprises primary (61%) and metastatic (39%) samples. This paper focuses on data from NSCLC patients in this cohort, which contains 1,494 metastatic and 1,855 primary samples. The gene panel has 481 genomic features.

The POPLAR (ClinicalTrials.gov ID: NCT01903993)^25^ and OAK (ClinicalTrials.gov ID: NCT02008227)^32^ clinical trials are, respectfully, phase 2 and phase 3 studies measuring the effectiveness of atezolizumab against docetaxel as a second line of treatment for patients with failed first-line platinum chemotherapy. The POPLAR study was conducted in Europe, Asia, and North America, and the OAK study was conducted in Europe, Asia, North America, South America, and New Zealand. Both studies enrolled adults (≥18 years old) with measurable disease per Response Evaluation Criteria in Solid Tumors (RECIST) version 1.1 and an Eastern Cooperative Oncology Group performance status of 0 or 1. Patients in both studies had previously received one or two cytotoxic chemotherapy regimens for stage IIIB or IV NSCLC, and those with *EGFR* mutations or *ALK* fusion oncogenes received tyrosine kinase inhibitor treatment.^33^ To identify the molecular mechanisms associated with immunotherapy, all analyses were conducted in the atezolizumab arm, which consists of 102 samples from POPLAR and 324 samples from OAK. The overlapped gene panels between OAK and POPLAR consists of 355 genomic features (Extended Data Fig. 4).

### Knowledge graphs: BIKG and Hetionet

The BIKG is AstraZeneca’s internal KG.^6,10^ It leverages all available AstraZeneca internal and external biological data sets, such as genes, pathways, cell lines, assays, compounds, and diseases. It includes more than 500 million relationships from more than 166 million publications and more than 50 databases, licensed repositories, and internal data sets. Graph-based knowledge allows us to infer evidence from similar features when there are missing data and sparse knowledge of a biological system. The BIKG also captures multiple biological mechanisms, which aids in the exploration of nonlinear biology and allows causal inference.

Hetionet is an integrative biomedical knowledge network^7^ that combines information from 29 public databases. The network contains 47,031 nodes of 11 types and 2,250,197 relationships of 24 types. Hetionet captures the conceptual distinctions between various components and mechanisms, such as genes and diseases or upregulation and binding.

### KG subgraphs for IO-related cancer treatment

To overcome the sparse molecular and missing biological information from the patient input data set, we constructed several BIKG subgraphs by integrating molecular and pathway biological information (Table 6). First, the following gene sets were identified:

**Immune-related gene set:** The latest version of a public database of innate immune interactions (InnateDB)^34,35^ was used to define an immune-relevant gene set (1,069 genes) for subgraph construction and context specificity. The gene-pathway mapping for this subset of genes was extracted from the baseline BIKG.
**CIViC disease specificity gene set:** CIViC (Clinical Interpretation of Variants in Cancer) was queried to identify 440 genes annotated as clinically relevant in NSCLC. The gene-pathway mapping for this subset of genes was extracted from the baseline BIKG.
**Biorelate NSCLC causal data:** Biorelate is a UK-based start-up company specializing in natural language processing–derived causal relationships between biomedical entities.
AstraZeneca licensed this data to extract approximately 108,000 gene-gene relationships specific to NSCLC (specifically, lung adenocarcinoma [LUAD] and lung squamous-cell carcinoma [LUSC]) based on literature evidence. All Biorelate data can be accessed via the company’s portal (https://www.biorelate.com).

**Table 6.** KG subgraphs evaluated in application to NSCLC, including IO related.

| **Subgraph** | **Description** | **No. of nodes** | **No. of edges** |
| --- | --- | --- | --- |
| 1 | Immune-related gene set with associated pathways | 23,915 | 1,297,468 |
| 2 | Immune-related gene set + CIViC NSCLC genes with associated pathways | 24,717 | 1,716,072 |
| 3 | Immune-related gene set + CIViC NSCLC genes | 20,134 | 1,524,005 |
| 4 | Biorelate NSCLC causal subgraph | 2,710 | 35,288 |
| 5 | Biorelate LUAD causal subgraph | 1,777 | 13,373 |
| 6 | CIViC NSCLC genes with associated pathways | 22,195 | 508,648 |
| Whole graph | Baseline BIKG graph | 27,349 | 4,185,751 |

Using these specific gene sets and their interactions reflected in BIKG, we then constructed subgraphs in which each type of subgraph provided context-specific (e.g., cancer type, patient subpopulation, mutation profile) querying and embedding methods (Table 6). The results in this manuscript are reported using a Biorelate NSCLC causal subgraph, which showed a consistent predictive performance and better stability of the results in the preliminary studies (Extended Data Table 1).

### KG feature extraction via graph embeddings for survival modeling

Creating structured, low-dimensional, patient-level embedding from unstructured graph data was critical for the next step, in which ML models were used to predict OS (Fig. 1b). The workflow consisted of three steps: (a) identifying patient-specific features in the graph through the patient genomic profile, (b) creating a gene-specific embedding from the graph, and (c) generalizing it to patient-level embedding.

1. From the gene panel, patient-level information was projected to the KG, exciting nodes, and associated pathways for each mutated gene in the patients’ profiles. The information was propagated through the graph via connected pathways.
2. Several methods were used to create gene-specific, fixed, low-dimensionality embedding from the KG, including GLEE,^17^ NetMF,^18^ RandNE,^19^ NodeSketch,^20^ BoostNE,^21^ and SocioDim.^16^ The SocioDim algorithm is based on identifying the largest eigenvalues from the gene-specific modularity matrix created from a patient-specific graph network. The process of learning gene latent representation from a fused graph of patient-gene and gene-gene interaction includes several steps. Given a patient heterogeneous graph *G*, we first calculated the *G*’s modularity matrix *B*:

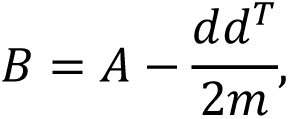

where *A* is the interaction matrix, *d* is the degree of each node, *d^T^* is the transpose matrix of *d*, and *m* is the number of edges in the graph *G*.

We then extracted the eigenvectors corresponding to the largest eigenvalues of the graph modularity matrix. The Lanzcos method^36^ was applied to calculate the top eigenvectors used as the gene embedding representation.

Patient embeddings were generated by averaging gene embeddings over all genes from the panel. We calculated the patient embedding as follows:

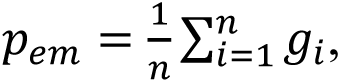

where *g_i_* is gene embedding and *n* is the number of gene mutations a patient carries.

### Survival modeling

We used Cox regression^37^ and random survival forest^38^ to model OS. Preliminary analysis demonstrated a much improved OS predictive performance with random survival forest, a nonparametric, nonlinear approach, so the results are reported here using this algorithm unless stated otherwise.

We compared input feature representations consisting of the gene panel with that from KG-based embedding derived using the gene panel to evaluate the importance of the KG in improving prediction of patient OS. For the IO-treated cohort, the results were also compared with the performance using TMB, a U.S. Food and Drug Administration–approved biomarker with a 10 Mut/Mb cutoff.^39^ In additional analyses, we evaluated the differences in the models’ predictive power by comparing primary with metastatic tissues.

### Training and evaluation approach

Data was split into training, test, and evaluation subsets. Bootstrapping was used for stability and sensitivity to training data. We used C-index for performance evaluation and KM survival analysis^23^ to stratify patients into high- and low-risk groups, inversely proportional to OS. We considered several cutoffs; results are reported here with the 75th percentile cutoff, which is widely used in the literature.^40–42^ The cutoff was determined by using training data during model development and applied to the evaluation data set.

### Biological interpretability

#### Key molecular features associated with prediction of patient survival

For each developed model, we identified the top 10 crucial gene features from the panel that contributed most to the OS prediction, using a decision tree regression model that associates embedding features with genomic features. We report the most important genomic features with a count of how many times they appear as important out of 10 independent models.

#### Cluster map to visualize top gene profiles

We plotted patients’ top gene profiles as a hierarchically clustered heat map to identify clusters of similar groups of mutations.

#### Differential analysis

The average value of mutation incidence was calculated for each gene. We compared patients’ high- and low-predicted risk groups to identify mutations associated with the high-risk group.

## Data availability

The pipeline will be publicly available soon.

Publicly available KGs that can be incorporated in the pipeline are PrimeKG,^5^ publicly available at https://github.com/mims-harvard/PrimeKG; and Hetionet,^7^ publicly available at https://het.io/.

## Acknowledgments

We thank Ben Sidders and Paul Metcalfe for their helpful suggestions, which improved the manuscript, the R&D IT BIKG team for their help with the BIKG data, and D.J. Shuman for editing help.

## Author contributions

[TODO: Add]

