## Supplemental data for "Integrating Knowledge Graphs into Machine Learning Models for Survival Prediction and Biomarker Discovery in Patients with Non–Small-Cell Lung Cancer"

**Extended Data Table 1 | Survival prediction comparing the performance of BIKG and Hetionet, using different subgraphs extracted from BIKG and their effects on OS prediction**

| | MSK (*N* = 1,494) | | | | OAK (*N* = 324) | | | |
|---|---|---|---|---|---|---|---|---|
| | BIKG | | Hetionet | | BIKG | | Hetionet | |
| **Subgraph** | **C-index** | ***P*** | **C-index** | ***P*** | **C-index** | ***P*** | **C-index** | **P** |
| Genomic features | 0.557 ± 0.026 | — | 0.557 ± 0.026 | — | 0.591 ± 0.052 | — | 0.591 ± 0.052 | — |
| Immune-relevant gene set with associated pathways | 0.559 ± 0.025 | 0.863 | 0.554 ± 0.028 | 0.087 | 0.589 ± 0.037 | 0.922 | 0.617 ± 0.066 | 0.341 |
| Immune-relevant gene set + CIViC NSCLC genes with associated pathways | 0.558 ± 0.025 | 0.931 | 0.551 ± 0.034 | 0.663 | 0.597 ± 0.055 | 0.805 | 0.601 ± 0.064 | 0.706 |
| Immune-relevant gene set + CIViC NSCLC genes | 0.552 ± 0.017 | 0.618 | 0.560 ± 0.032 | 0.821 | 0.592 ± 0.066 | 0.97 | 0.602 ± 0.065 | 0.681 |
| Subgraph consists of CIViC NSCLC genes with associated pathways | 0.563 ± 0.027 | 0.619 | 0.572 ± 0.031 | 0.257 | 0.599 ± 0.059 | 0.751 | 0.628 ± 0.070 | 0.198 |
| Whole KG | 0.547 ± 0.014 | 0.303 | 0.552 ± 0.034 | 0.716 | 0.599 ± 0.052 | 0.735 | 0.603 ± 0.061 | 0.642 |
| Biorelate NSCLC + LUAD + LUSC contexts causal subgraph | 0.562 ± 0.031 | 0.701 | N/A | N/A | 0.621 ± 0.051 | 0.209 | N/A | N/A |
| Biorelate LUAD contexts causal subgraph | 0.560 ± 0.031 | 0.817 | N/A | N/A | 0.611 ± 0.061 | 0.441 | N/A | N/A |

BIKG, Biological Insights Knowledge Graph; C-index; concordance index; CIViC, Clinical Interpretation of Variants in Cancer; KG, knowledge graph; LUAD, lung adenocarcinoma; LUSC, lung squamous-cell carcinoma; MSK, Memorial Sloan Kettering Cancer Center; N/A, not applicable; NSCLC, non–small-cell lung cancer; OS, overall survival.

**Extended Data Table 2 | OncoKG[a] for top genes identified in the OAK and POPLAR data sets**

| Gene | Therapeutic level | Diagnostic level | Prognostic level | Resistance level | FDA level | Alterations | Reference | Comments |
|---|---|---|---|---|---|---|---|---|
| *TP53* | | | Px1 | | | Oncogenic | | |
| *EGFR* | 1 | | | R1 | 2 | Oncogenic | | |
| *ATM* | 1 | | | | 2 | Oncogenic | | |
| *KRAS* | 1 | Dx2 | | R1 | 2 | Oncogenic | | |
| *EPHA7* | | | | | | Likely oncogenic | Yu et al.[1] | Very little data |
| *STAT3* | | Dx3 | | | | Oncogenic | Zheng et al.[2] | Claims to overcome EGFRi resistance |
| *CTNNB1* | | | | | | Oncogenic | Muto et al.[3] | |
| *PRKDC* | | | | | | | Pan et al.[4] | Internal interest |
| *NFE2L2* | | | | | | Oncogenic | Zhang et al.[5] | |
| *SOX9* | | | | | | Oncogenic | Grimm et al.[6] | |

[a]OncoKB is a precision oncology knowledge base developed at Memorial Sloan Kettering Cancer Center that contains biological and clinical information about genomic alterations in cancer.

EGFRi, epidermal growth factor receptor inhibitor.

**Extended Data Table 3 | OncoKG[a] for top genes identified in the MSK data set**

| Gene | Therapeutic level | Diagnostic level | Prognostic level | Resistance level | FDA level | Alterations | Reference |
|---|---|---|---|---|---|---|---|
| *STK11* | 4 | | | | | Oncogenic | Di Federico et al.[7] |
| *KRAS* | 1 | Dx2 | | R1 | 2 | Oncogenic | |
| *KEAP1* | | | | | | Oncogenic | Di Federico et al.[7] |
| *TP53* | | | Px1 | | | Oncogenic | |
| *SMARCA4* | | | | | | Oncogenic | Zhou et al.[8] |
| *ATM* | 1 | | | | 2 | Oncogenic | Chen et al.[9] |
| *PTPRD* | | | | | | Oncogenic | Wang et al.[10] |
| *EPHA3* | | | | | | Likely oncogenic | Bai et al.[11] |
| *RBM10* | | | | | | Oncogenic | Zhang F et al.[12] |
| *FAT1* | | | | | | Oncogenic | Zhang W et al. 2022[13] |

[a]OncoKB is a precision oncology knowledge base developed at Memorial Sloan Kettering Cancer Center that contains biological and clinical information about genomic alterations in cancer.

**Extended Data Table 4 |Average incidence of gene mutations compared between high- and low-risk groups for OS prediction models trained with a, BIKG, and b, OAK gene panel. .**

| Patient risk level | *TP53* | *EGFR* | *ATM* | *KRAS* | *EPHA7* | *STAT3* | *CTNNB1* | *PRKDC* | *NFE2L2* | *SOX9* |
|---|---|---|---|---|---|---|---|---|---|---|
| | **BIKG using OAK gene panel** | | | | | | | | | |
| High | 0.961 | 0.153 | 0.076 | 0.038 | 0.077 | 0.000 | 0.000 | 0.077 | 0.154 | 0.000 |
| Low | 0.394 | 0.092 | 0.171 | 0.092 | 0.053 | 0.014 | 0 | 0.052 | 0.092 | 0.000 |
| **OAK gene panel** | | | | | | | | | | |
| High | 0.769 | 0.038 | 0.038 | 0.192 | 0.038 | 0.000 | 0.000 | 0.153 | 0.115 | 0.000 |
| Low | 0.461 | 0.132 | 0.184 | 0.039 | 0.066 | 0.013 | 0.000 | 0.026 | 0.105 | 0.000 |

**Extended Data Table 5 | Average incidence of gene mutations compared between high- and low-risk groups for OS prediction models trained with a, BIKG, and b, MSK gene panel.**

| Patient risk level | *STK11* | *KRAS* | *KEAP1* | *TP53* | *SMARCA4* | *ATM* | *PTPRD* | *EPHA3* | *RBM10* | *FAT1* |
|---|---|---|---|---|---|---|---|---|---|---|
| **BIKG using MSK gene panel** | | | | | | | | | | |
| High | 0.422 | 0.467 | 0.467 | 0.578 | 0.111 | 0.044 | 0.044 | 0.066 | 0.044 | 0.089 |
| Low | 0.053 | 0.303 | 0.068 | 0.378 | 0.045 | 0.045 | 0.083 | 0.083 | 0.121 | 0.053 |
| **MSK gene panel** | | | | | | | | | | |
| High | 0.444 | 0.400 | 0.644 | 0.400 | 0.200 | 0.089 | 0.156 | 0.111 | 0.133 | 0.156 |
| Low | 0.045 | 0.325 | 0.007 | 0.439 | 0.015 | 0.030 | 0.045 | 0.068 | 0.091 | 0.030 |

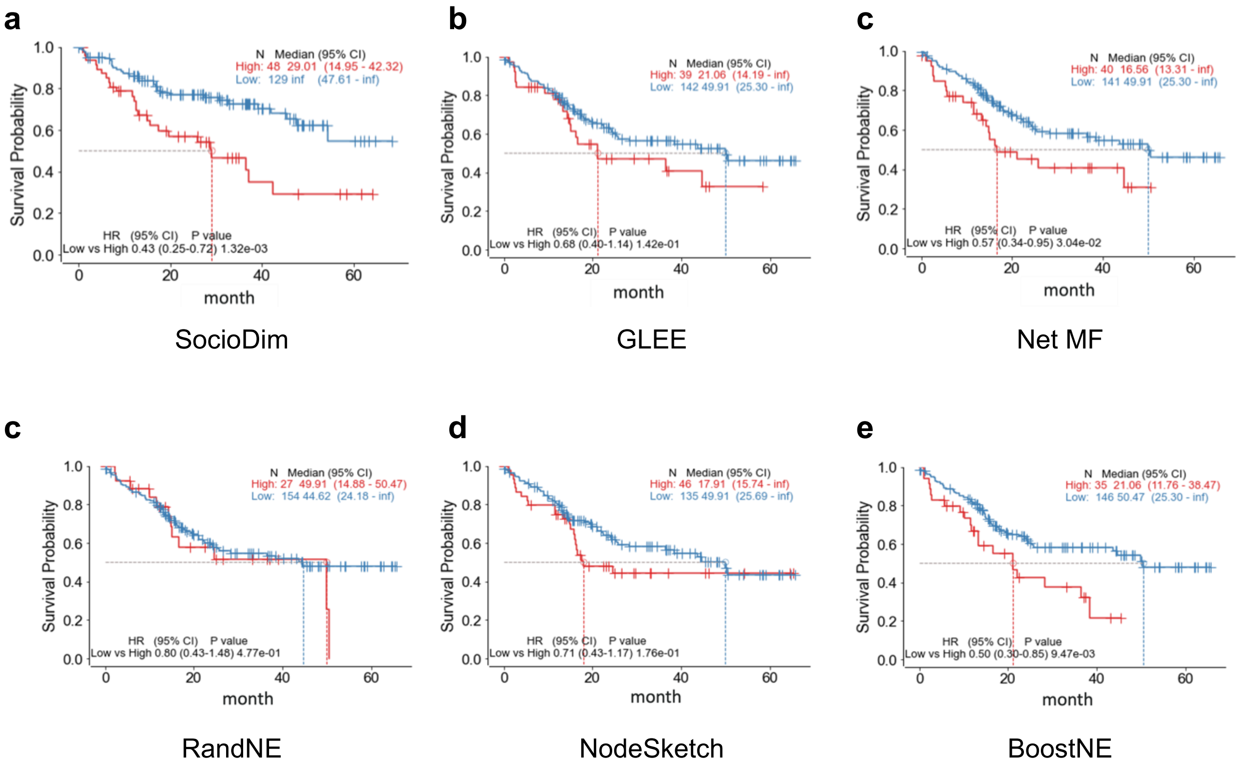


**Extended Data Fig. 1 | Comparison of different graph embedding approaches.** With models trained on MSK, evaluation results are shown using MSK holdout sets.

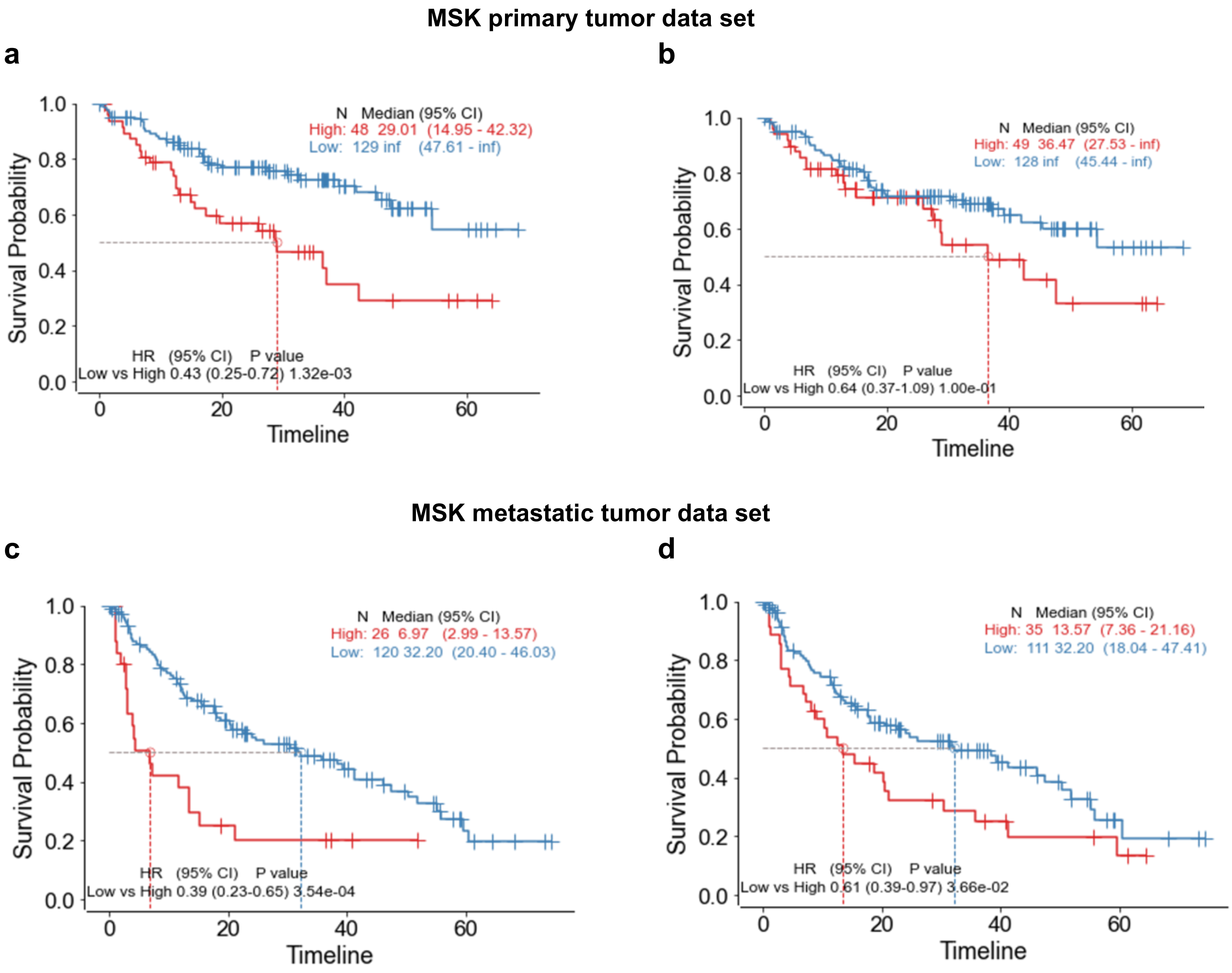


**Extended Data Figure 2 | Use of BIKG prior knowledge to stratify patient survival on MSK primary and metastatic tumor sample data sets.** **a**, **b**, Patient survival stratification using the MSK primary tumor sample data set. **a**, Survival prediction model trained with BIKG prior knowledge; **b**, survival prediction model trained using MSK gene panel features. **c**, **d**, Patient survival stratification using the MSK metastatic tissue tumor sample data set. **c**, Survival prediction model trained with BIKG prior knowledge; **d**, survival prediction model trained with the MSK gene panel. Survival prediction models trained with BIKG prior knowledge resulted in statistically significant patient stratification in both the MSK primary and metastatic tumor sample data sets.

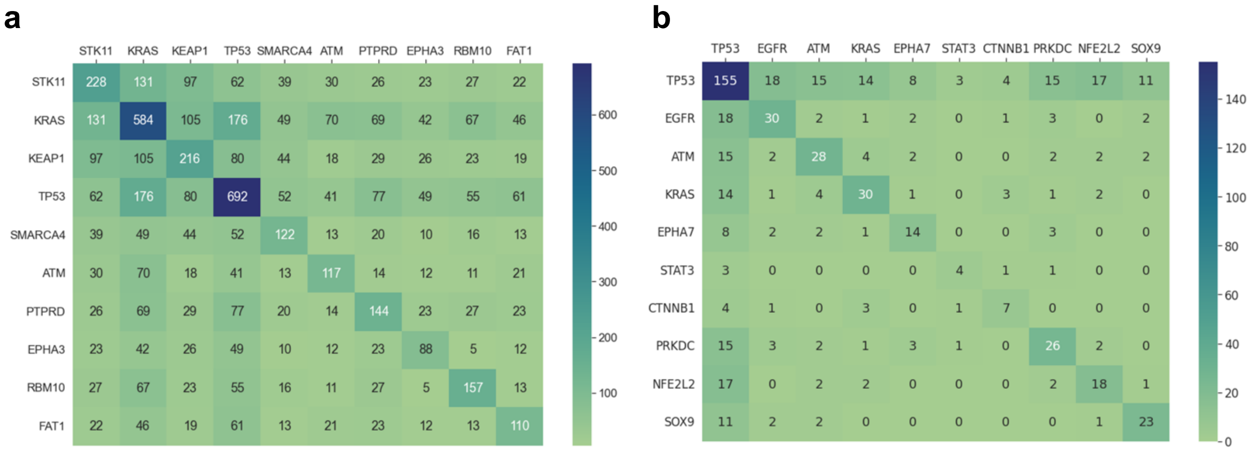


**Extended Data Fig. 3 | Gene co-occurrence matrix of data sets. a**, MSK data set; **b**, OAK-POPLAR data set. Rows and columns show the top 10 genes, and values in the matrices represent the number of co-occurrences between two genes. Shading indicates the most commonly co-occurring genes in each panel.

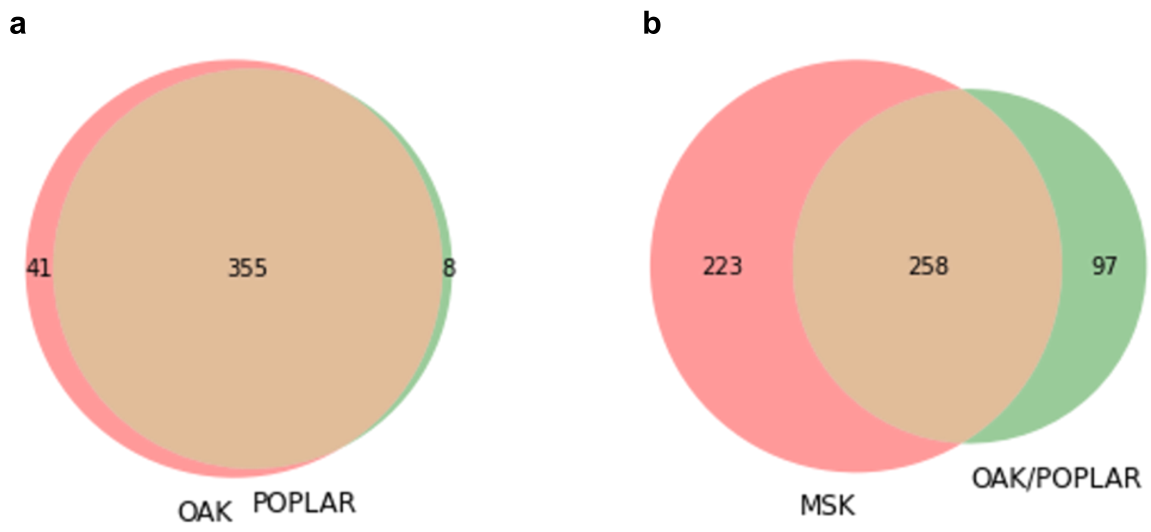


**Extended Data Fig. 4 | Venn diagram of gene panels and their overlap. a**, Overlap between the OAK and POPLAR gene panels. A total of 355 genes overlapped between the two panels. **b**, Overlap between MSK and OAK-POPLAR gene panel (i.e., the intersection between OAK and POPLAR). The MSK and OAK-POPLAR panels have 258 genes in common. The MSK panel has 223 genes that are excluded from the OAK-POPLAR panel, and the OAK-POPLAR panel has 97 genes that are excluded from the MSK panel.
